# Geny: A Genotyping Tool for Allelic Decomposition of Killer Cell Immunoglobulin-Like Receptor Genes

**DOI:** 10.1101/2024.02.27.582413

**Authors:** Qinghui Zhou, Mazyar Ghezelji, Ananth Hari, Michael K.B. Ford, Connor Holley, COVNET Consortium, Lisa Mirabello, Stephen Chanock, S. Cenk Sahinalp, Ibrahim Numanagić

## Abstract

Accurate genotyping of Killer cell Immunoglobulin-like Receptor (KIR) genes plays a pivotal role in enhancing our understanding of innate immune responses, disease correlations, and the advancement of personalized medicine. However, due to the high variability of the KIR region and high level of sequence similarity among different KIR genes, the currently available genotyping methods are unable to accurately infer copy numbers, genotypes and haplotypes of individual KIR genes from next-generation sequencing data. Here we introduce Geny, a new computational tool for precise genotyping of KIR genes. Geny utilizes available KIR haplotype databases and proposes a novel combination of expectation-maximization filtering schemes and integer linear programming-based combinatorial optimization models to resolve ambiguous reads, provide accurate copy number estimation and estimate the haplotype of each copy for the genes within the KIR region. We evaluated Geny on a large set of simulated short-read datasets covering the known validated KIR region assemblies and a set of Illumina short-read samples sequenced from 25 validated samples from the Human Pangenome Reference Consortium collection and showed that it outperforms the existing genotyping tools in terms of accuracy, precision and recall. We envision Geny becoming a valuable resource for understanding immune system response and consequently advancing the field of patient-centric medicine.

## 1 Introduction

The natural killer (NK) cells are a critical component of the human innate immune system, which is the first line of host defense mechanisms against infections, viruses, and diseases. These cells are responsible for rapid response to various pathological challenges, such as viral-infected cells and cancerous cells [1–3]. The NK cells are regulated by cell surface receptors that interact with major histocompatibility complex class I (MHC-I) molecules found on the surface of various cells in the body [4]. These receptors are, in turn, encoded by Killer cell Immunoglobulin-like Receptor (KIR) genes, located on the human chromosome 19 within a 150kb region of the Leukocyte Receptor Complex (LRC), whose expression and interactions are essential for distinguishing between healthy and abnormal cells.

The KIR genes contribute to the wide array of immune responses observed among individuals due to their vast genetic diversity which also influences disease susceptibility [5]. For that reason, KIR genes belong to the family of *highly polymorphic genes* and consequently harbor a myriad of known haplotypes (also known as *alleles*, or in some cases *genotypes*) that are present among the human population [6]. Importantly, this variation is not limited only to the coding regions; it also encompasses the regulatory regions that direct the expression of KIR genes. It has been proposed that this vast genetic diversity likely stems from the evolutionary pressures posed by constantly evolving viruses [7]. Such intricate genetic architecture means that fewer than 2% of unrelated individuals share an identical KIR genotype [8].

The seventeen (17) KIR genes are named based on their extracellular Immunoglobulin-like (lg-like) domains (designated as 2D or 3D) and the lengths of their cytoplasmic tails (marked as L for long cytoplasmic tails, S for short cytoplasmic tails, and P for pseudogene). A general rule is that short-tailed KIRs are activating receptors, while long-tailed KIRs are inhibitory receptors. Based on these designations, the KIR genes can be categorized as follows: (a) six (6) genes with two domains and long cytoplasmic tails (*KIR2DL1* – *KIR2DL5B*), (b) five (5) genes with two domains and a short cytoplasmic tail (*KIR2DS1* –*KIR2DS5*), (c) three (3) genes with three domains and long tails (*KIR3DL1* –*KIR3DL3*), (d) one (1) *KIR3DS1* that is characterized by having three domains and a short tail, and (e) two (2) pseudogenes (*KIR2DP1* and *KIR3DP1*) 1. The whole-region KIR haplotypes are divided into two categories: group B (having one of *KIR2DL5, KIR2DS1, KIR2DS2, KIR2DS3, KIR2DS5* and *KIR3DS1*) and group A (having none of these genes) [7]. Finally, names of individual gene haplotypes, or alleles, roughly follow the star-allele nomenclature introduced for annotating pharmacogenes [9], where each allele is assigned a number that indicates its function [8]. The current known KIR haplotypes (alleles) have been assembled and cataloged within the IPD-KIR database [10].

As different KIR alleles result in different immune responses, it is necessary to precisely genotype and haplotype KIR genes in order to better understand the role these genes play within the immune system. One cost-effective way of doing that is by using high-throughput sequencing (HTS) technologies that have been successfully used for large-scale genotyping [11]. However, KIR genotyping cannot be easily done through the established HTS genotyping pipelines, such as GATK [12], primarily due to the high gene polymorphism of individual KIR genes. Not only KIR genes harbor many variants, but their alleles are defined by the whole gene haplotype—resolving this haplotype necessitates both variant calling and phasing. Another reason is that the copy number of each KIR gene varies significantly across individuals: while the presence of some genes is relatively uncommon (e.g., *KIR2DS3*), it is not rare to see some genes with large copy numbers (e.g., *KIR2DL4* or *KIR3DP1*), where each copy may have a different allele. Finally, the sequence contents of many KIR genes are mutually similar, which introduces high levels of ambiguity during the alignment of short reads to the KIR region. Such ambiguity is typically resolved in an arbitrary fashion, which produces incorrect alignments and, in turn, incorrect variant and allele calls. All these challenges are exacerbated by the reference genome itself: the latest canonical version of the human genome (GRCh38) does not include most of the KIR genes in the primary assembly and has no consistent reference model of the whole KIR region.

Some of these challenges have been previously encountered and addressed within the context of pharmacogene genotyping [13–15]. Genes such as *CYP2D6, CYP2A6, CYP2C19* and *SLCO1B1*, also exhibit high levels of polymorphism and are subject to various copy number and structural variation events, which makes them incompatible with the standard genotyping pipelines. Thus, many specialized genotyping tools specifically tailored for pharmacogenes have been recently proposed. Of these tools, Aldy [16], Cypiripi [17], PyPGx [18], StellarPGx [19], Stargazer [20] and Astrolabe [21] stand out as premier genotyping approaches for pharmacogenes. However, despite their success in the field of pharmacogenomics, these tools rely on the correct and precise alignments to the target genes to make correct allele calls and are unable to handle complex regions such as KIR, where most of the read alignments are ambiguous. While one of these tools, Aldy 4 [22], provides some support for reads alignment within the *CYP2D* region, it cannot handle the scale and complexity of 17 KIR genes.

One genomic region that shares similar ambiguous alignment problems as KIR but has been successfully genotyped is the immunoglobulin heavy chain locus. The variable genes (IGHV) present in this locus are particularly challenging to genotype, with high polymorphism rate, copy number variants, structural variants, and homologous sequences [23]. This problem has been addressed by the ImmunoTyper-SR tool [24, 25], which uses a combinatorial optimization approach to resolve read mapping and alignment ambiguities. However, while IGHV genes are numerous (*∼* 120 functional and non-functional copies per chromosome), they are much shorter than KIR genes (*∼* 280 bp vs 13.4 kbp), and the resulting difference in scale means that this approach cannot be utilized for KIR genotyping.

For these reasons, a few tools have been recently developed to assess the KIR region itself from sequencing data. These include T1K [26], PING [27], KASS [28], KPI [29] and KIR*IMP [30]. Some tools, such as KPI, only handle gene-level identification and are unable to precisely call individual KIR alleles. KASS relies on *de novo* assembly of error-corrected sequences from PacBio’s long-read capture data that are annotated with KIR genes and exon/intron locations. Finally, tools such as PING, KIR*IMP, and T1K can identify individual alleles from the short-read sequencing data. PING utilizes *k*-mer fingerprinting to call individual alleles but is hard to run as it requires manual parameter estimation for each input cohort, and it also overlooks specific genes that are highly similar to each other, such as *KIR3DL1* and *KIR3DS1* [26]. Another approach, KIR*IMP, relies on a statistical SNP imputation to call KIR haplotypes but is limited to highquality SNPs and sufficiently large reference panels. Finally, T1K utilizes an expectation maximization strategy to rapidly identify KIR and HLA (Human Leukocyte Antigen) genotypes from sequencing data. While T1K offers speed and acceptable accuracy, it currently lacks the capability to determine the copy number of KIR genes. It is also worth noting that many of these tools call alleles based solely on their sequence similarity to the reference KIR alleles and thus sometimes fail to distinguish alleles by their true functional impact, as sequence similarity is not a perfect proxy for functional characterization of sequences.

In order to address the outstanding challenges in analyzing and genotyping the KIR region, we introduce Geny, a GENotYper for KIR genes. This tool combines an expectation minimization-based filtering scheme with a combinatorial optimization approach in the form of integer linear programming (inspired by our previous pharmacogenomics tools Aldy [16]) to infer copy number and the exact allele of each present KIR gene copy. Furthermore, it can detect and leverage all variant types found in the KIR database and is able to distinguish between core, allele-defining variants that define the allele’s functionality and the silent variants that have no major impact on the overall functionality. We show that Geny achieves better precision and recall—up to 20%—over the existing KIR callers on both simulated and real datasets and that it provides significantly fewer miscalls than the other tools. Finally, to demonstrate a real-world application of Geny, we generated KIR genotypes on a 461 patients COVID-19 cohort from the COVNET Consortium ^1^ and perform statistical association analysis between KIR alleles and COVID-19 severity. As such, we hope that Geny lays the groundwork for precise KIR genotyping algorithms and that it will become a major part of future biomedical applications dealing with human immune system behavior.

## 2 Results

We assessed the performance of Geny and other major tools, T1K and PING, on two large datasets: simulated reads on top of fifty (50) completely assembled KIR regions from GenBank and on 25 whole genome Illumina-sequenced HPRC samples. We also use Geny to genotype KIR genes on 419 patients from the COVID-19 cohort from the COVNET Consortium and to infer associations between the various KIR alleles and the COVID-19 severity.

During the assessment of the performance of each tool, we computed the number of differences between the ground truth call and the inferred call, where the number of misses corresponds to the number of false positives and false negatives. We also provide the standard precision, recall and F1 scores for each tool. Note that we limited ourselves only to functional allele concordance (i.e., the first three-digit match of the allele name; thus, allele *\*0010101* is treated as *\*001*). While Geny calls the complete allele name, other tools, such as T1K, only provide the functional portion of the allele call. Furthermore, we observed that many alleles in our datasets were novel and did not exactly match any of the alleles in the database, mostly due to the differing silent variants, and thus did not have an established name. Finally, note that some tools, such as PING, may output multiple possible solutions. In these cases, we selected the allele option that is closest to the ground truth and reported those as representative calls.

We compared Geny’s calls with T1K and PING, the only comparable KIR genotyping tools that provide allele-level genotype calls. However, we encounter several challenges when attempting to apply PING. Firstly, PING assumed samples from the same cohort. We also note that running PING required a lot of manual intervention and manual parameter inference, as the default set of parameters produced suboptimal results (see Supplementary Materials for details); on the other hand, both Geny and T1K required only input FASTQ or SAM/BAM/CRAM files to operate. PING also seemed to be extremely sensitive to the user-provided probe hit ratio thresholds for setting copy numbers of each gene. Finally, PING assumes that only one copy of *KIR3DL3* is present per haplotype to normalize the number of *k*-mer hits per gene in their copy number estimation stage. While they suggest using *KIR3DL2* to normalize the *k*-mer hit counts in case *KIR3DL3* is duplicated in a sample [31], it is unclear how to do so. As our simulated dataset did not satisfy the first and third criteria (cohort data and fixed *KIR3DL3* copy number), we were not able to apply PING to this dataset.

### 2.1 Ground truth annotations

The ground truth for each sample was obtained by analyzing the complete assemblies of the KIR region. The annotations were generated using the KIR-Annotator tool which was developed specifically for this application. Initially, the KIR allele database was aligned to the assembly with minimap2 [32], followed by the merging of all overlapping mappings to locate putative genes. The gene type was identified by selecting the gene with the highest number of alleles mapped. Subsequently, the wildtype of the identified gene was re-mapped to the putative gene sequence to refine its location again using minimap2. The refined putative gene sequence was then aligned with the wildtype sequence via global alignment using parasail [33], allowing for the calling of variants and identification of functional variants. The closest allele to the putative gene sequence was selected based on the allele with the lowest functional variant Jaccard distance relative to the wildtype sequence, employing non-functional Jaccard distance in cases of ties. In other words, we prefer alleles that preserve their functionality by (1) having all its core variants present (see Methods for the exact definitions) and (2) not introducing novel core variants. In the case of a tie, the allele with the smallest Jaccard distance from the wildtype sequence was selected.

In some instances, the second condition could not be fulfilled without breaking the first condition. Even if both conditions are satisfied, the assembled sequence might still differ from the KIR-IPD allele sequences due to the differences in silent variations. Both cases point out that the sequenced allele is novel and is not yet cataloged within the IPD-KIR database; in either case, we selected the database allele that is closest to the observed allele as the “ground truth” based on the above criteria.

Finally, we performed some manual interventions on top of KIR-Annotator calls. In the case of GenBank assemblies, we used the existing GenBank allele annotations where possible to cross-validate and correct our calls. We also manually checked the presence of exon 1 deletion within *KIR3DP1* region that KIR-Annotator was unable to detect on its own.

### 2.2 Simulated data

We collected 50 complete assemblies of the KIR region from the GenBank [34], each corresponding to a distinct individual (Supplementary Materials). These assemblies cover a diverse set of KIR configurations, including cases with copy number variations, alleles from haplotype classes A and B, non-identified alleles and so on. Once these sequences were annotated, each was independently inserted within an assembly of chromosome 19(chr19:54,724,235–54,867,216) to replace the KIR locus and create a synthetic KIR assembly sample. We then simulated perfect paired-end reads of size 100bp that cover this locus with the coverage of 20*×* for each synthetic assembly. In order to create diploid samples, representing the 2 copies of the KIR locus present in a human genome, we randomly selected pairs of synthetic assemblies and combined their simulated reads to create 21 synthetic diploid samples. The resulting samples encompassed all 15 KIR genes, the two pseudogenes and contain 828 true alleles spread across these genes. The allele count for each gene is shown in Table 1.

**Table 1:** Comparison of Geny and T1K on simulated datasets from the 50 GenBank assemblies. Bold type indicates better results. Geny produces a significantly lower number of miscalls and outperforms T1K in all metrics, in some cases by a large margin (up to 20%).

| Gene | Total | Geny |  |  |  | T1K |  |  |  |
| --- | --- | --- | --- | --- | --- | --- | --- | --- | --- |
|  |  | Misses | Precision | Recall | F1 | Misses | Precision | Recall | F1 |
| *KIR2DL1* | 66 | **4** | **98.4%** | **95.4%** | **0.97** | 10 | 91.0% | 93.8% | 0.92 |
| *KIR2DL2* | 29 | **5** | **89.7%** | **92.9%** | **0.91** | 6 | 89.3% | 89.3% | 0.89 |
| *KIR2DL3* | 55 | **0** | **100.0%** | **100.0%** | **1.00** | 5 | 96.3% | 94.5% | 0.95 |
| *KIR2DL4* | 81 | **0** | **100.0%** | **100.0%** | **1.00** | 5 | **100.0%** | 93.8% | 0.97 |
| *KIR2DL5A* | 23 | **2** | **92.0%** | **100.0%** | **0.96** | 8 | 75.9% | 95.7% | 0.85 |
| *KIR2DL5B* | 13 | **4** | **100.0%** | 69.2% | **0.82** | 70 | 13.6% | **100.0%** | 0.24 |
| *KIR2DP1* | 68 | **5** | **94.0%** | **98.4%** | **0.96** | 9 | 91.0% | 95.3% | 0.93 |
| *KIR2DS1* | 26 | **3** | **96.0%** | **92.3%** | **0.94** | 4 | 95.8% | 88.5% | 0.92 |
| *KIR2DS2* | 28 | **4** | **87.1%** | **100.0%** | **0.93** | 8 | 80.0% | 92.3% | 0.86 |
| *KIR2DS3* | 16 | **0** | **100.0%** | **100.0%** | **1.00** | 40 | 27.8% | 93.8% | 0.43 |
| *KIR2DS4* | 62 | **2** | **98.4%** | **98.4%** | **0.98** | 14 | 83.3% | 96.8% | 0.90 |
| *KIR2DS5* | 18 | **0** | **100.0%** | **100.0%** | **1.00** | **0** | **100.0%** | **100.0%** | **1.00** |
| *KIR3DL1* | 61 | **4** | **100.0%** | **93.4%** | **0.97** | 11 | 88.5% | 93.1% | 0.91 |
| *KIR3DL2* | 87 | **7** | **96.4%** | 95.2% | **0.96** | 33 | 69.7% | **95.8%** | 0.81 |
| *KIR3DL3* | 82 | **11** | **91.0%** | 94.7% | **0.93** | 33 | 68.0% | **98.6%** | 0.80 |
| *KIR3DP1* | 89 | **14** | **90.7%** | **92.9%** | **0.92** | 26 | 84.1% | 86.0% | 0.85 |
| *KIR3DS1* | 24 | **0** | **100.0%** | **100.0%** | **1.00** | 6 | 87.0% | 87.0% | 0.87 |
| All | 828 | **65** | **95.8%** | **96.1%** | **0.96** | 288 | 75.5% | 93.6% | 0.84 |

Geny has more than 200 fewer misses over T1K, as can be seen in the results shown in Table 1. It also improves the precision by 20% and F1 score by 0.12. Geny outperformed T1K on all individual genes as well. We note that T1K had a high false positive rate on *KIR2DL5B* and *KIR2DS3* (71 and 40, respectively). Also, it struggled with the KIR3DL gene family. However, its recall was competitively high, albeit slightly lower than Geny’s. We also note that Geny also had issues with the KIR3DL family, particularly with *KIR3DP1* and *KIR3DL3*, where it often completely missed the presence of these genes or exonic deletions that define some of the core *KIR3DP1* variants.

The cases where Geny misses the allele can be roughly explained as follows: (1) novel or non-standard alleles that have a non-standard combination of core variants or large exonic deletions and, as such, get filtered out; (2) an “extended” solution where the true allele mistakenly gets assigned an additional core variant due to incorrectly resolved cross-gene read alignments; and (3) copy number inconsistencies. While we plan to address cases (1) and (3) in the near future, we note that the second case is challenging to handle because the wrong solution can be explained by the observable reads based on the current model.

### 2.3 Real Data

To evaluate the performance of Geny on real data, we conducted a comparative analysis using 25 whole genome samples sequenced by Illumina NovaSeq 6000 (read length 150bp) sourced from the 1000 Genomes Project [35]. The ground truth for this comparison was derived from multi-model assemblies generated by the Human Pan Genome Consortium [36] and covers diverse ethnicities. As such, this dataset ensures that the evaluation reflects a highly realistic assessment of tools’ performance in real-world scenarios.

As can be seen in Table 2, Geny again outperformed all other tools with respect to all metrics (precision, recall, F1 score and miss rate). For example, Geny misses nearly two times fewer alleles than T1K and more than three times as compared to PING. In terms of precision, recall and F1 score, Geny again outperforms other tools. It also performs well on individual KIR genes. The sole exceptions are the *KIR3DL2* and *KIR3DL3* genes, where PING produces the best results. In general, it is *KIR3DL3* and *KIR3DP1* that cause most of the trouble for T1K and Geny on this dataset; this is not surprising, as *KIR3DP1* has already been reported to pose significant challenges for correct genotyping [27]. In general, Geny’s misses follow the same patterns as observed on the simulation datasets. We note that many assemblies point to the existence of novel and uncatalogued KIR alleles; further work will be necessary to validate and catalog them correctly.

**Table 2:**
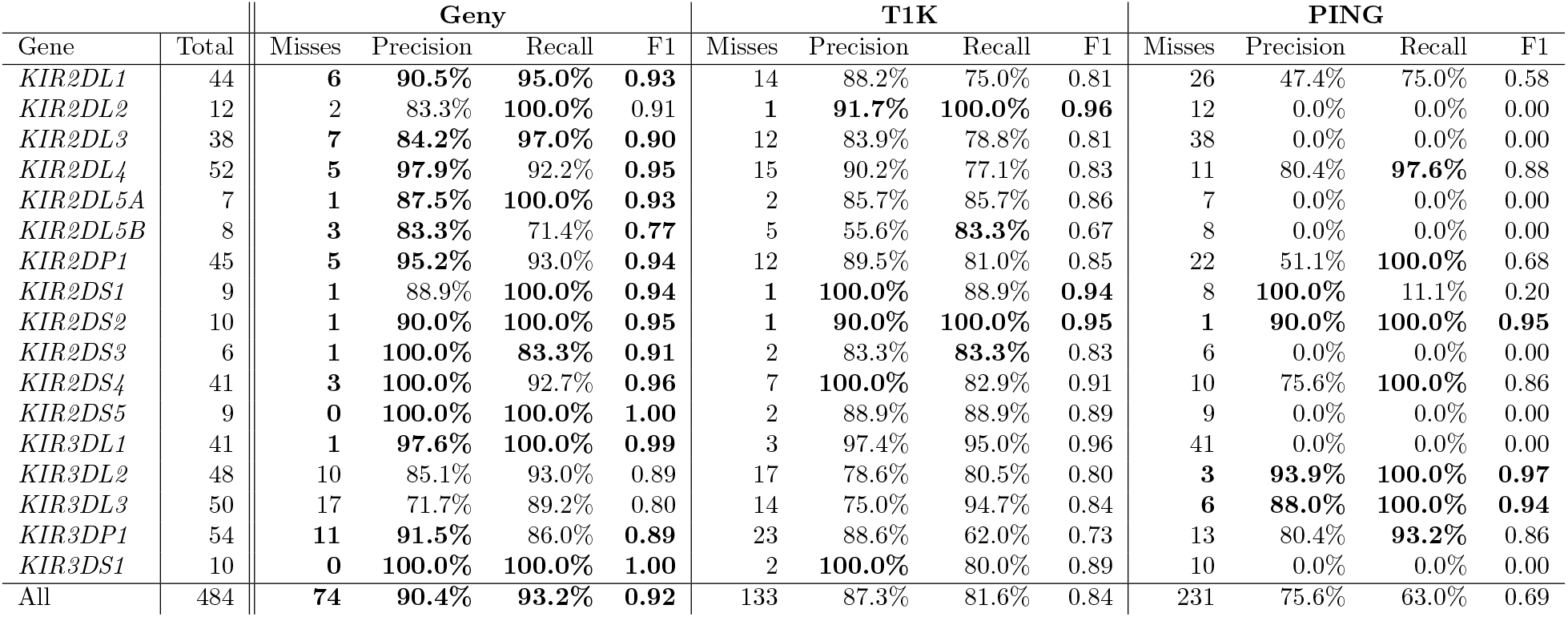
Comparison of Geny, PING and T1K on 25 HPRC samples. Bold type indicates better results. Geny again outperforms other tools by a large margin in many of the metrics. The exceptions are the *KIR3DL2–3* genes, on which both Geny and T1K underperform when compared to PING.

**Table 3:**
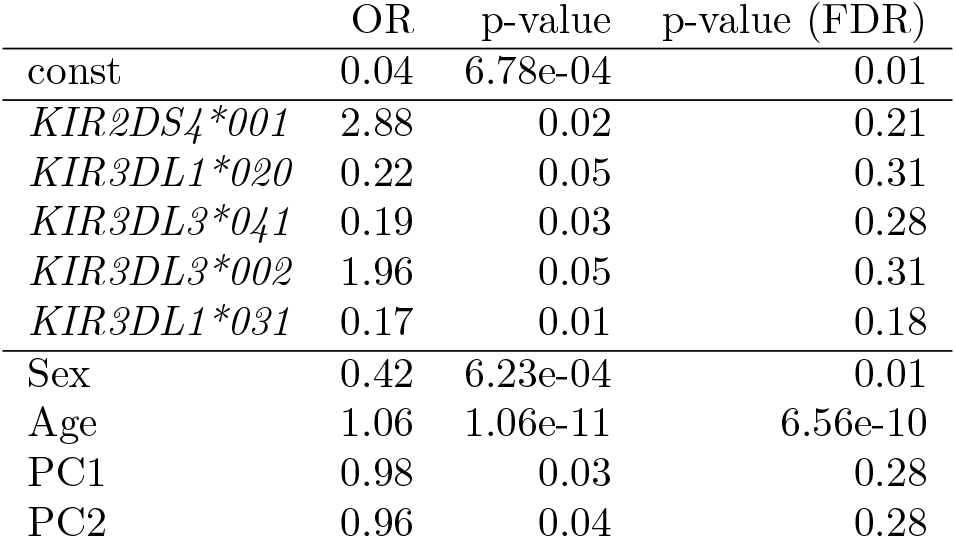
Results from COVID association for all independent variables and covariates with p *≤* 0.05 before FDR correction. OR = Odds ratio

|  | OR | p-value | p-value (FDR) |
| --- | --- | --- | --- |
| const | 0.04 | 6.78e-04 | 0.01 |
| *KIR2DS4\*001* | 2.88 | 0.02 | 0.21 |
| *KIR3DL1\*020* | 0.22 | 0.05 | 0.31 |
| *KIR3DL3\*041* | 0.19 | 0.03 | 0.28 |
| *KIR3DL3\*002* | 1.96 | 0.05 | 0.31 |
| *KIR3DL1\*031* | 0.17 | 0.01 | 0.18 |
| Sex | 0.42 | 6.23e-04 | 0.01 |
| Age | 1.06 | 1.06e-11 | 6.56e-10 |
| PC1 | 0.98 | 0.03 | 0.28 |
| PC2 | 0.96 | 0.04 | 0.28 |

On a final note, the current version of Geny typically takes from a few minutes to around half an hour on more complex samples to genotype all genes within a sample. PING takes, on average, around half an hour per sample, while T1K typically finishes in under a minute.

### 2.4 COVID-19 Associations

For the preliminary use of an allele-level whole-genome sequencing (WGS) KIR genotyping tool in disease association studies, we applied Geny to a set of 461 WGS samples from COVID-19 patients generated by the COVNET Consortium. COVID-19 severity was classified as mild (cases with no hospitalization, death, or respiratory support due to COVID-19), moderate (cases with hospitalization for COVID-19, no death, and non-mechanical respiratory support), or severe (cases with death or hospitalization for COVID-19 and mechanical respiratory support), and this information was binarized by merging moderate and severe cases into a single category. Samples with insufficient or unclear data were excluded, as were samples with inconsistent sex data. This resulted in 285 and 176 samples with mild and moderate/severe COVID-19 disease, respectively, totaling 461 samples. Genetic principal components, previously generated by the COVNET Consortium using the WGS data, were utilized as covariates for population stratification control. The genotype data were refined by first binarizing the presence of KIR functional alleles and then filtering out alleles that are present in less than 5% of the samples and those with the perfect mutual correlation or high multicollinearity via variance inflation factor calculation (threshold = 10). The remaining alleles were confirmed to lack multicollinearity by calculating condition indices (maximum value = 8.4). The sample size was assessed in relation to the number of remaining independent variables, which included 56 KIR allele presence/absence binary variables and covariates of binarized sex, age, and the top 3 principal components, with a ratio of 2.88 samples to the smallest dependent variable category.

The resulting logistic regression model, incorporating 53 KIR allele variables along with the covariates, was fitted using `statsmodels` [37]. The model’s fit was evaluated using Pseudo-R-Squared (value of 0.23) and the Hosmer-Lemeshow test (*p* = 1.0), which suggested a reasonable fit and good model performance. We found 5 alleles significantly associated with COVID-19 severity, exhibiting *p*-values below 0.05. Of these 5 alleles, 3 belong to genes *KIR2DS4* and *KIR3DL1*, which have previously been associated with COVID-19 disease severity [38].

However, after applying false discovery rate (FDR) correction for multiple hypothesis test correction, no KIR allele associations remained significant, while the sex and age covariates remained significant after correction. We also repeated this analysis using complete gene presence (instead of allele-level presence) as independent variables, but again, no significant associations were found after all necessary corrections. Despite the good fit and performance indicated by all evaluation metrics, we note that the ratio of sample size to independent variables fell below the recommended threshold of 10, hinting at a potential for overfitting and underscoring the necessity for further validation within a larger cohort.

## 3 Discussion

The process of genotyping and haplotyping KIR genes is important for a deeper understanding of the innate immune system and its interactions. Here we have presented Geny, a new tool for identification of KIR alleles within high-throughput sequencing datasets. In our evaluations, Geny consistently outperformed the current state-of-the-art methods for KIR genotyping across many metrics on a diverse set of samples. As we move towards tailored medical treatments, the accuracy of tools like Geny in identifying genes can shape the future of patient care.

Despite its performance, there are still many areas left for improvement and further study. One of them is the detection of novel major alleles—functional alleles that have not been cataloged by the existing KIR databases. This also includes calling of fusion alleles that have been observed in the wild [39]. Another room for improvement is the quality of ground truth data. Currently, we have no externally validated set of ground truth calls for KIR genes as we have in pharmacogenomics (e.g., GeT-RM project [40]). We hope that the recent availability of HPRC samples and assemblies will result in this issue being resolved soon. Finally, we note that many parts of the Geny pipeline can be optimized in terms of speed; for that purpose, we plan to rewrite the current Python implementation in a high-performance language such as C++ or Codon [41].

## 4 Methods

### 4.1 Overview

The Geny pipeline consists of three major stages. The first stage reads the short-read HTS data from a FASTQ or a SAM/BAM file and computes all possible alignments to the reference KIR sequences for each read found in the sample. The second stage filters out unlikely KIR alleles and reads assignments by employing both deterministic and statistical criteria to reduce the overall search space and enhance the quality of the final calls. Finally, the third stage solves the integer linear programming (ILP) model that determines the correct copy number and the exact haplotypes of each present KIR gene.

### 4.2 Preliminaries: Notation and Database Preparation

Each Geny stage requires the annotated KIR haplotype database. We use the latest version of the IPD-KIR database [10] (v2.12.0) that contains the haplotype sequences of currently known KIR alleles, as well as the associated allele names that characterize their functionality. Each name is a set of at most seven (7) digits, where the first three digits indicate the allele functionality (typically defined by the non-synonymous exonic changes), the next two digits indicate the synonymous exonic changes, and the last two digits indicate all other changes [2]. In the rest of the section, we will utilize the terminology from pharmacogenomics [22] and will refer to alleles with different functionality as *major alleles*. For example, *KIR2DL2*0010101* and *KIR2DL2*0010105* both encode same major allele (*KIR2DL2*001*) and thus the same protein, while only differing by a couple of non-exonic variants.

Determination of the functional behavior of present KIR alleles (in other words, *major allele* calling) is the key aspect of the KIR genotyping process. The functional behavior is, in turn, defined by the set of *core variants*: variants that distinguish the functionality of a given allele from the other alleles. While these variants are typically functional (including both SNPs and indels of various sizes), they can also include UTR variants, whole exon deletions and other variants that affect gene expression. All other variants that do not impact the allele’s function are called *silent variants*.

Unlike pharmacogenomics databases such as PharmVar, the KIR-IPD database contains only the haplotype sequences for each allele and does not provide a list of core variants that differentiates those alleles from the reference (wildtype) allele (typically denoted as *\*001* or *\*0010101*). Most of the available annotation tools, even when annotating complete assemblies, only rely on a simple edit distance score to compare allele sequences and oftentimes fail at properly determining the correct allele calls because they do not distinguish between core (functional) and silent variants. For example, *KIR2DS1*011* allele is defined by the core c.5812 G*>*A functional variant that distinguishes its functionality from *KIR2DS1*002* (the reference allele). While many other variants also distinguish *\*011*’s sequence from *\*002*’s, they are either silent or intronic and can be ignored when testing for the presence of *\*011*. However, if all variants are considered the same (as they are in edit distance calculation), the lack of a few silent variants will overcome the concordance of a single core variant and might result in the wrong major allele assignment. To avoid this issue, we developed a PharmVar-like allele database for each KIR gene by aligning each allele sequence from the KIR-IPD database to the reference allele with parasail [33] and calculating the list of core variants that define each allele. We also established the complete genomic sequence for each allele: while the IPD-KIR database contains complete sequences for most of its alleles, there are cases where it only provides the coding sequence or a small exonic part that differentiates the allele from the reference allele. Finally, based on the existing literature (e.g., [42, 43]) and GenBank annotations, we constructed a KIR locus reference sequence that contains all 17 KIR genes and used it during the alignment step.

Formally, the final Geny database contains a set of KIR genes *𝒢* = *{𝒢*_1_, …, *𝒢*_17_*}*. Each gene *𝒢*_*g*_ *∈ 𝒢* harbors a list of variants *ℳ*_*g*_ = *{m*_*g*,1_, *m*_*g*,2_, … *}* and a set of alleles *𝒜*_*g*_ = *{𝒜*_*g*,1_, *𝒜*_*g*,2_, … *}*. The allele *𝒜*_*g*,1_ is considered to be *reference allele*. Let *𝒜* = ⋃*g 𝒜*_*g*_. Each allele *𝒜*_*g*,*i*_ is in turn defined by a variants *ℳ*_*g*,*i*_ *⊆ ℳ*_*g*_. Each variant *m*_*g*,*j*_ *∈ ℳ*_*g*_ is a tuple (*l*_*g*,*j*_, *o*_*g*,*j*_) containing its location *l*_*g*,*j*_ in the reference allele *𝒜*_*g*,1_ and an operation *o*_*g*,*j*_ (SNP or an indel). For example, the previously mentioned c.5812 G*>*A is encoded as (5812, *GA*). A mutation *m*_*g*,*j*_ *∈ ℳ*_*g*_ is a core variant iff core(*m*_*g*,*j*_) = 1. A location *l* in gene *𝒢*_*g*_ is called *core location* if there is allele *𝒜*_*g*,*i*_ *∈ 𝒜*_*g*_ that has a core variant at location *l*. Finally, each allele *𝒜*_*g*,*i*_ is assigned **a**_*g*,*i*_ that corresponds to its genomic sequence. **a**_*g*,*i*_[*l*] indicates the *l*-th position in such sequence.

### 4.3 Stage 1: Alignment

The first step of Geny pipeline aligns the input reads *ℛ* = *{r*_1_, *r*_2_, … *}* to the allele sequences **a**_*g*,*i*_. Because many reads in the KIR region can be aligned to many different alleles across many genes, Geny needs to compute all alignments from each read to each KIR allele from the database. We use minimap2 [32] in all-to-all mode (`--dual=no -P --secondary=yes`) to achieve this. Following the alignment, Geny discards all alignments that contradict the core variants for each allele, severely clipped alignments and those that have a low alignment score. Finally, we end up with a set of alignments 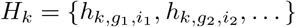 for each read *r*_*k*_ *∈ ℛ*. Each alignment indicates the target allele sequence **a**_*g*,*i*_, as well as the location on it and the edit operation needed to align the read to it.

In order to be able to determine the copy number of each KIR gene, we also align input reads to a copy number-neutral region in the genome. By default, we use copy number-neutral *COMT* gene region; other choices can be provided by the end user. Alignment to the copy number-neutral region provides the expected coverage of the sequencing data that is used later to call copy numbers and alleles.

### 4.4 Stage 2: Filtering

The large number of the KIR alleles—the current version of the KIR-IPD database contains more than 1,500 known alleles among 17 KIR genes—adversely impacts the search space of the subsequent combinatorial optimization step. Therefore, Geny attempts to limit the number of valid alleles by filtering out those that are unlikely to occur based on the alignment data.

In the first pass, Geny selects only those alleles whose core variants are covered by a sufficient number of reads are considered. By default, we set the minimum allowed read coverage to 3. We also need to ensure that alleles that do not harbor a core variant at a core variant site still have sufficient coverage at that site to be considered.

#### 4.4.1 Landmark generation

In addition to filtering out the unlikely alleles, it is also important to filter out the spurious read alignments. Ideally, we would like to consider only a small set of alignments that map to the core locations of each allele remaining after the previous filtering and discard other reads. However, as many reads that map to the core locations also map to regions within other genes that contain no core locations specific to that gene, we need to extend the set of core locations to also include other locations that “mirror” the core locations in other genes. Thus, we introduce the concept of *landmark locations* that are projections of the valid core locations to all alleles (of any gene) that, despite not harboring core variants, still “catch” the reads that cover core variants in other alleles. The objective of landmarks is to provide the opportunity for the reads to be assigned to non-variant harboring alleles.

To infer landmarks, we construct an overlap graph *G*_*g*,*i*_ for each candidate allele *𝒜* _*g*,*i*_ *∈ 𝒜*_*g*_ to capture the relationships between the alignments that cover the variants *ℳ*_*g*,*i*_ (Figure 2). We consider all alignments *h*_*k*,*g*,*i*_ which either cover a core location or for which there is another alignment *h*_*k*,*g*_*′*,*i′* in another gene or allele that covers a core location. Each such alignment *h*_*k*,*g*,*i*_ corresponds to a node in the graph. A graph edge is created between two alignments if they overlap on **a**_*g*,*i*_. Constructing the overlap graph enables us to identify the *landmark regions* in **a**_*g*,*i*_ that harbor “interesting” reads; those regions can be found by finding the strongly connected components (SCCs) within each *G*_*g*,*i*_. Once landmark regions are identified, we establish landmark locations *L*_*g*_ = *l*_1_, *l*_2_, … for each gene *𝒢*_*g*_ by augmenting the set of gene’s core locations with the minimal number of other positions in *𝒢*_*g*_ so that each alignment that covers a landmark region also covers at least one landmark position. Finally, we select all alignments that are covered by a landmark position and discard the others.

**Figure 1:**
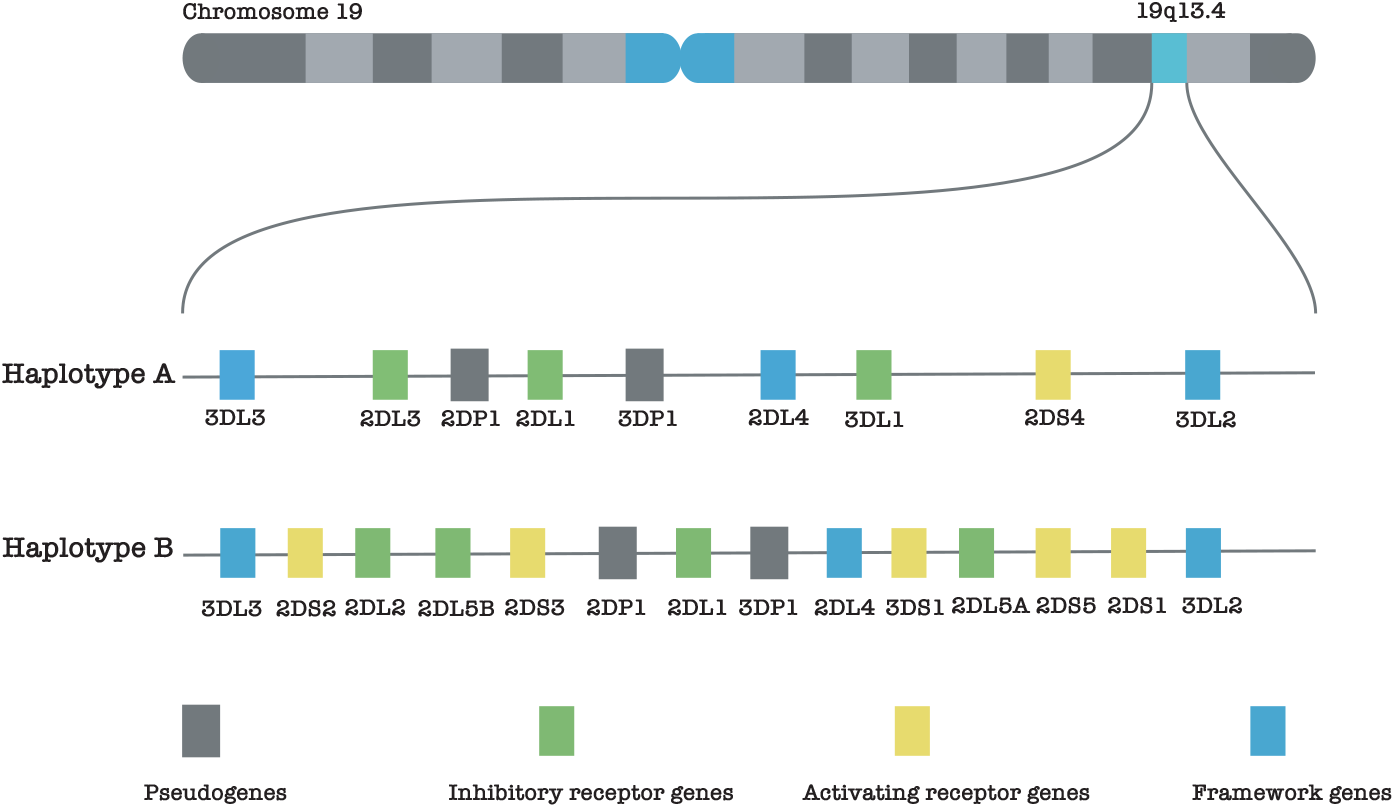
Illustration of the KIR gene positions on chromosome 19, showcasing the distinct structures of haplotype groups A and B.

**Figure 2:**
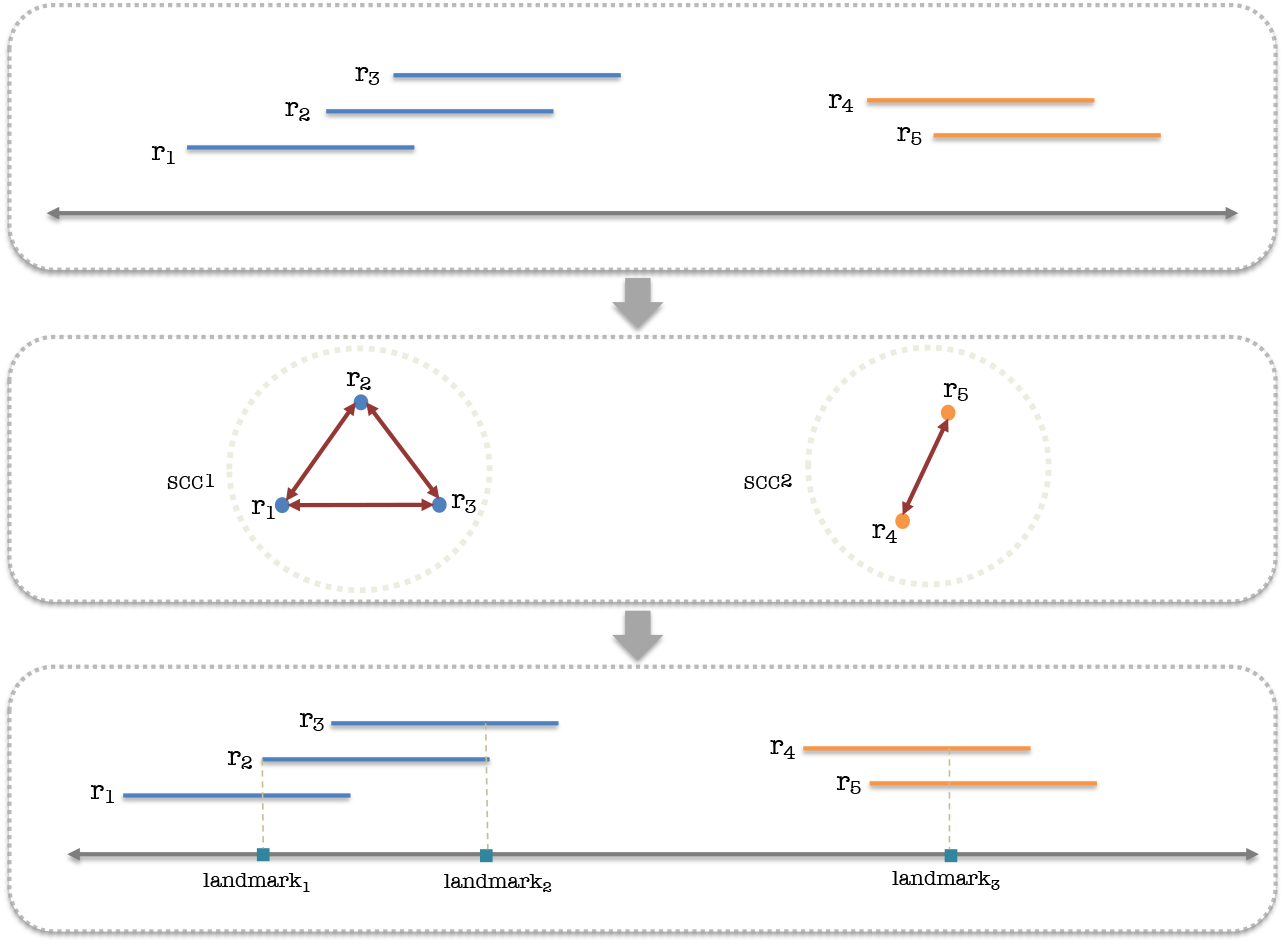
Diagram illustrating the process of landmark generation. Initially, all valid reads are collected as input. These reads are then utilized to construct a graph that represents their overlap. Then, we identify strongly connected components (SCCs) of that graph, which represent groups of reads that continually cover a region by overlapping each other. In the concluding step, an iterative approach is used to determine the minimal set of landmarks needed to capture all the valid reads within each SCC. With each iteration, the number of landmarks is increased until all reads in a given SCC are captured.

#### 4.4.2 Candidate allele selection

After obtaining a set of valid alleles and reads, Geny further filters the set of valid alleles through the Expectation Maximization (EM) algorithm [44] by identifying alleles with lower densities in the input sample, thus reducing the solution space for the final solver and improving specificity [45]. The EM algorithm, in a setting where the input data is partially known and the parameters of the distribution function (model) that generated the data are unknown, iteratively estimates the parameters of the model to maximize the likelihood of the observed data. We perform maximum likelihood estimation on the abundance of each allele and the sequencing error rate in a similar fashion as in [46].

Let *ϕ* denote the abundance of each of the *n* = | *𝒜*| candidate alleles, *ℒ* (*θ*) the log-likelihood of the read set *ℛ* consisting of *m* = | *ℛ*| total reads given the parameter *θ, Ƶ*_*k*_ the latent variable representing the allele which generates *r*_*k*_, and *ϕ*_*i*_ the density of allele *𝒜*_*i*_ *∈ 𝒜*. Let the sequencing error rate be *ϵ*. Consider there are 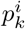 matching bases for mapping read *r*_*k*_ *∈ ℛ* on *𝒜*_*i*_ *∈ 𝒜* with allele length *l*_*i*_. To account for multiple alignments, we denote 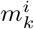 to be the number of positions read *r*_*k*_ maps on allele *𝒜*_*i*_. Then:

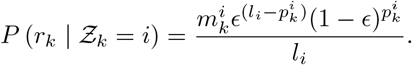

We define the log-likelihood *ℒ* (*θ*) as follows:

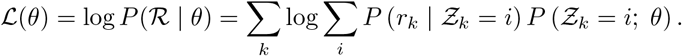

Following this, we obtain the EM update steps for parameters *ϕ* and *ϵ* as follows (see Supplementary Materials for details). Let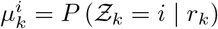. Then:

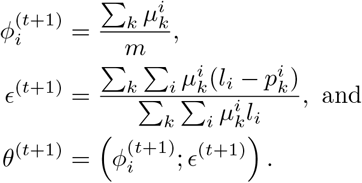

Once the updated parameter values are obtained, we select all alleles associated with components of non-trivial presence, where *ϕ > γ*, for further refinement in the final stage and discard the others. Unlike other methods that use EM to select the final solution, we set *γ* to a small value and only use this step *to filter out unlikely candidates*. This ensures that Geny avoids the common problem with EM-based methods, where the final solution ends up being a local maximum that is not relevant to the true call.

### 4.5 Stage 3: Allele Calling

For the final phase of allele calling, we aim to apply the Integer Linear Programming model (ILP) to assign each read to a proper KIR allele that passed the previous filtering stages and select the true alleles present in the sample. The problem is set as follows.

For each read *r*_*k*_ *∈ ℛ* and allele *𝒜*_*g*,*i*_ *∈ 𝒜*, we introduce a binary variable *V*_*k*,*g*,*i*_ that is set if and only if *h*_*k*,*g*,*i*_ is the best alignment among all candidate alignments *H*_*k*_^2^. In other words, *V*_*k*,*g*,*i*_ indicates that the read *r*_*k*_ is assigned to *𝒜*_*g*,*i*_. We also allow the possibility of dropping reads—i.e., not assigning it to either of the alleles—by introducing the variable *D*_*k*_ = 1 *− Σg*,*i V*_*k*,*g*,*i*_. Let us also introduce an integer variable *A*_*g*,*i*_ that is set to the number of times allele *𝒜*_*g*,*i*_ is selected in the final solution. Let the constant *ξ* denote the expected coverage of a single allele copy (determined via copy number-neutral region in Stage 1).

An integer linear program that selects a set of alleles among *𝒜* and assigns the reads to them so as to minimize the difference between the observed coverage and the selected one can be formulated as follows:

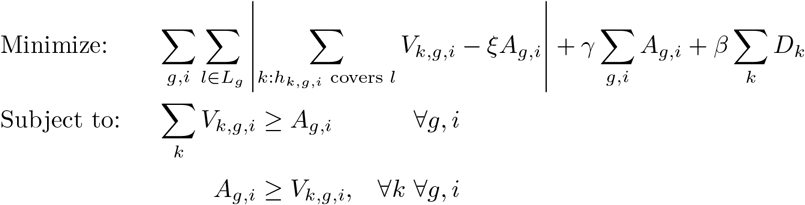

*L*_*g*_ is a set of landmarks for the gene *G*. A selection cost *γ* is associated with each selected allele, and a read drop cost *β* for discarding each read.

By formulating the problem as an ILP and minimizing the total absolute coverage error, we effectively optimize the assignment of reads to alleles, resulting in more accurate and reliable allele identification. We employed Gurobi as a reliable tool for solving ILP problems in an efficient manner [47]. Note that, unlike models used in pharmacogenonics (e.g., Aldy 4 [22]), this model performs read selection and is thus an order of magnitude more complex than the previous models. Currently, the model deploys tens of thousands of binary variables and hundreds of continuous variables.

## Software availability

Geny is available at https://github.com/0xTCG/geny and also uploaded as a Supplemental Code. The experimental procedure and results are available at https://github.com/0xTCG/geny/tree/master/paper and are also uploaded as Supplemental Notebook and Supplemental Experiments, respectively.

## Competing interests statement

None declared.

## Acknowledgements

We recognize the valuable contributions made by these collaborators to the COVNET Consortium:

Evangelos Andreakos Laboratory of Immunobiology, Center for Clinical, Experimental Surgery and Translational Research, Biomedical Research Foundation of the Academy of Athens, Athens, Greece

Mary Carrington Basic Science Program, Frederick National Laboratory for Cancer Research, NCI, Frederick, MD USA

Laboratory of Integrative Cancer Immunology, Center for Cancer Research, National Cancer Institute, Bethesda, MD USA Ragon Institute of MGH, MIT and Harvard, Cambridge, MA USA

Robert J. Kreitman Laboratory of Molecular Biology, Center for Cancer Research, National Cancer Institute, Bethesda, MD USA Sudha Sharma Department of Biochemistry and Molecular Biology, National Human Genome Center,

Howard University College of Medicine, Washington, DC USA Mark S. Brahier Georgetown University School of Medicine, Washington, DC USA

Heather S. Feigelson Institute for Health Research, Kaiser Permanente Colorado, Aurora, CO USA

Rex L. Chisholm Center for Genetic Medicine, Northwestern University Feinberg School of Medicine, Chicago, IL USA

Hogune Im Genome Opinion, Inc., Seoul, Republic of Korea

Bruce R. Korf Department of Genetics, University of Alabama at Birmingham, Birmingham, AL USA

Marylyn D. Ritchie Department of Genetics, Perelman School of Medicine, University of Pennsylvania, Philadelphia, PA USA

Leslie G. Biesecker Center for Precision Health Research, National Human Genome Research Institute, Bethesda, MD USA

Clifton L. Dalgard Uniformed Services University of the Health Sciences, Bethesda, MD USA

Amy A. Hutchinson Cancer Genomics Research Laboratory, Frederick National Laboratory for Cancer Research, Frederick, MD USA

Xin Li Cancer Genomics Research Laboratory, Frederick National Laboratory for Cancer Research, Frederick, MD USA

Brooke Rosenblum Center for Precision Health Research, National Human Genome Research Institute, Bethesda, MD USA

Sharon A. Savage Division of Cancer Epidemiology and Genetics, National Cancer Institute, Rockville, MD USA

Vibha Vij Division of Cancer Epidemiology and Genetics, National Cancer Institute, Rockville, MD USA

Meredith Yeager Cancer Genomics Research Laboratory, Frederick National Laboratory for Cancer Research, Frederick, MD USA

Department of Biology, Hood College, MD, USA

Lisa Mirabello Division of Cancer Epidemiology and Genetics, National Cancer Institute, Rockville, MD USA

## Author Contributions

S.C.S, and I.N. designed the study. Q.Z. and M.G. developed the methods and the software, and conducted the experiments. M.F. and A.H. conducted the experiments and performed COVNET data analysis. L.M. and S.C. provided the COVNET data. C.H. and I.N. adapted the KIR database. M.G., Q.Z., A.H., M.F., S.C.S. and I.N. wrote the manuscript.

## Funding

Q.Z., M.G. and I.N. were supported by National Science and Engineering Council of Canada (NSERC) Discovery Grant (RGPIN-04973), Canada Research Chairs Program, Canada Foundation for Innovation’s John R. Evans Leaders Fund (CFI JELF) and B.C. Knowledge Development Fund (BCKDF). C.H. was supported by the BioTalent SWPP program. A.H., M.F. and S.C.S. were supported by funding from the Intramural Research Programs of the National Cancer Institute (NCI). A.H. is also funded by the NCI-UMD Partnership Program.

## Appendix A EM algorithm derivation

Given the likelihood function with respect to parameters *θ* as:

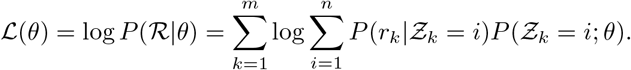

We assume each read *r*_*k*_ being uniformly sampled from each allele *𝒜*_*i*_ with length of allele *l*_*i*_, and each base of the read are independently generated with a sequencing error rate *ϵ*, and there are 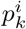 matching bases mapping *r*_*k*_ on *𝒜*_*i*_. Considering multi-mapping of read *r*_*k*_ within allele *𝒜*_*i*_, assume read *r*_*k*_ maps to 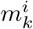 positions on allele *𝒜*_*i*_, we have:

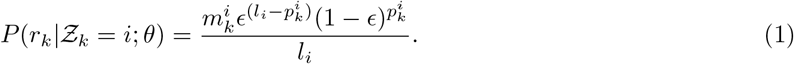

Let

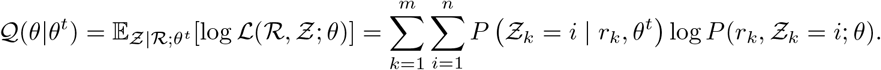

For each *i, j* we compute:

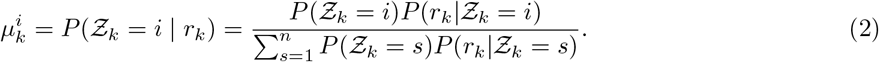

To maximize *Q*(*θ*|*θ*^*t*^), we construct a Lagrangian:

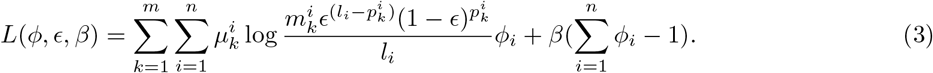

To update *ϵ*, by KKT condition, we set:

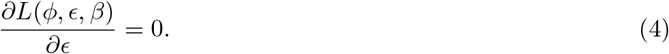

Then:

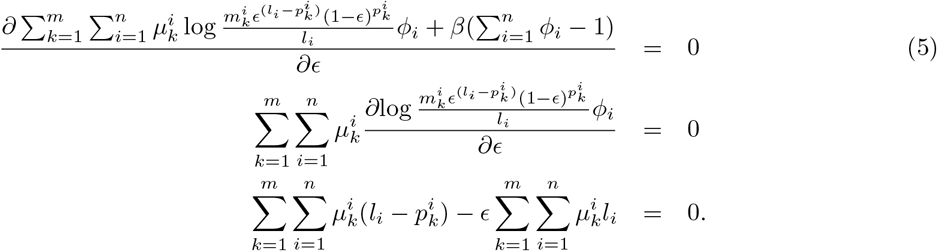

Thus:

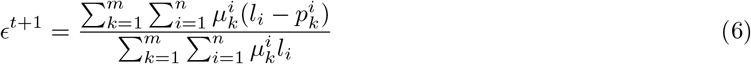

To update *ϕ*_*i*_, set:

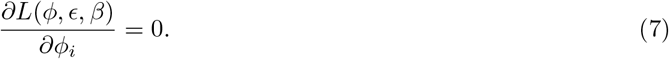

Thus:

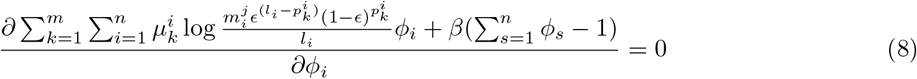

We have:

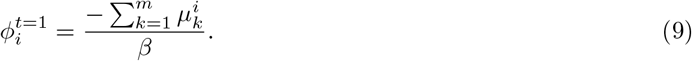

Since 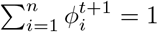 we have 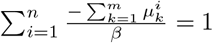 thus:

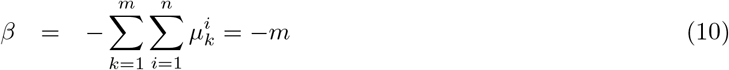

Applying (10) to (9), we get:

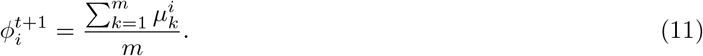

## Appendix B Accession Numbers

GenBank IDs that contain complete KIR region assemblies and were used for simulations:

- NT 113949.2
- NT 187636.1
- NT 187637.1
- NT 187638.1
- NT 187639.1
- NT 187640.1
- NT 187641.1
- NT 187642.1
- NT 187643.1
- NT 187644.1
- NT 187645.1
- NT 187668.1
- NT 187669.1
- NT 187670.1
- NT 187671.1
- NT 187672.1
- NT 187673.1
- NT 187674.1
- NT 187675.1
- NT 187676.1
- NT 187677.1
- NT 187683.1
- NT 187684.1
- NT 187685.1
- NT 187686.1
- NT 187687.1
- NT 187693.1
- NW 003571054.1
- NW 003571055.2
- NW 003571056.2
- NW 003571057.2
- NW 003571058.2
- NW 003571059.2
- NW 003571060.1
- NW 003571061.2
- NW 016107300.1
- NW 016107301.1
- NW 016107302.1
- NW 016107303.1
- NW 016107304.1
- NW 016107305.1
- NW 016107306.1
- NW 016107307.1
- NW 016107308.1
- NW 016107309.1
- NW 016107310.1
- NW 016107311.1
- NW 016107312.1
- NW 016107313.1
- NW 016107314.1

Accession numbers for 25 HPRC samples used in the experimental section:

- HG00438
- HG00673
- HG00735
- HG00741
- HG01071
- HG01106
- HG01175
- HG01258
- HG01358
- HG01361
- HG01891
- HG01928
- HG01952
- HG01978
- HG02148
- HG02257
- HG02572
- HG02622
- HG02630
- HG02717
- HG02886
- HG03453
- HG03516
- HG03540
- HG03579

## Appendix C Running PING on the Validation Dataset

We used the version of PING from https://github.com/wesleymarin/PING/tree/wgs_snakemake, as mentioned in [48], since we wanted to run it on WGS data. We extracted paired-end FASTQ files from the 25 HPRC WGS BAMs which were fed to PING. The tool was given 32 GB of memory and had access to 16 CPUs; scratch space was 100 GB.

We used the following probe hit ratio thresholds (Supplementary Table 1) for the copy numbers of each KIR gene, based on the initial plots generated by PING. The table is in the same format as the example threshold file (`manualCopyThresholds example.csv`) that was packaged with the tool.

**Supplementary Table 1:** Thresholds for probe hit ratios (mean number of probe hits per target KIR gene divided by the mean number of *KIR3DL3* probe hits) used to generate validation results on PING.

|  | 0-1 | 1-2 | 2-3 | 3-4 | 4-5 | 5-6 |
| --- | --- | --- | --- | --- | --- | --- |
| *KIR3DP1* | 0.15 | 0.35 | 0.6 | 0.75 | NA | NA |
| *KIR2DS5* | 0.2 | 0.65 | 1.1 | NA | NA | NA |
| *KIR2DL3* | 0.17 | 0.6 | 1 | NA | NA | NA |
| *KIR2DP1* | 0.3 | 0.6 | 1.2 | NA | NA | NA |
| *KIR2DS3* | 0.2 | 0.8 | 1.4 | NA | NA | NA |
| *KIR2DS2* | 0.2 | 0.5 | NA | NA | NA | NA |
| *KIR2DL4* | 0.2 | 0.5 | 1.2 | 1.6 | NA | NA |
| *KIR3DL3* | 0 | 0 | NA | NA | NA | NA |
| *KIR3DL1* | 0.2 | 0.7 | 1 | NA | NA | NA |
| *KIR3DS1* | 0.2 | 0.7 | 1 | NA | NA | NA |
| *KIR2DL2* | 0.15 | 0.35 | 0.65 | 1.1 | NA | NA |
| *KIR3DL2* | 0.2 | 1 | 1.7 | NA | NA | NA |
| *KIR2DS4* | 0.2 | 0.85 | NA | NA | NA | NA |
| *KIR2DL1* | 0.2 | 0.6 | 0.8 | 1.2 | NA | NA |
| *KIR2DS1* | 0.15 | 0.4 | NA | NA | NA | NA |
| *KIR2DL5* | 0.15 | 0.5 | 0.8 | 1.1 | NA | NA |

## Footnotes

1 https://dceg.cancer.gov/research/how-we-study/genomic-studies/covnet

2 For the sake of explanation and to avoid notation abuse, we assume that each read can only be aligned to a single location within a given allele; in practice, this is not true but the overall model applies to this case as well.

## Notes

### Competing Interest Statement

The authors have declared no competing interest.

